# Human immune and gut microbial parameters associated with inter-individual variations in COVID-19 mRNA vaccine-induced immunity

**DOI:** 10.1101/2022.08.08.503075

**Authors:** Masato Hirota, Miho Tamai, Sachie Yukawa, Naoyuki Taira, Melissa M. Matthews, Takeshi Toma, Yu Seto, Makiko Yoshida, Sakura Toguchi, Mio Miyagi, Tomoari Mori, Hiroaki Tomori, Osamu Tamai, Mitsuo Kina, Eishin Sakihara, Chiaki Yamashiro, Masatake Miyagi, Kentaro Tamaki, Matthias Wolf, Mary K. Collins, Hiroaki Kitano, Hiroki Ishikawa

## Abstract

COVID-19 mRNA vaccines induce protective adaptive immunity against SARS-CoV-2 in most individuals, but there is wide variation in levels of vaccine-induced antibody and T-cell responses. However, factors associated with this inter-individual variation remain unclear. Here, using a systems biology approach based on multi-omics analyses of human blood and stool samples, we find that baseline expression of AP-1 transcription factors, *FOS* and *ATF3*, is inversely correlated with BNT162b2 mRNA vaccine-induced T-cell responses. *FOS* expression is associated with transcription modules related to baseline immunity, but it is negatively associated with those related to T-cell activation upon BNT162b2 mRNA stimulation. Interestingly, the gut microbial fucose/rhamnose degradation pathway is positively correlated with *FOS* and *ATF3* expression and inversely correlated with BNT162b2-induced T-cell responses. Taken together, these results demonstrate that baseline expression of AP-1 genes, which is associated with the gut microbial fucose/rhamnose degradation pathway, is a key negative correlate of BNT162b2-induced T-cell responses.

## Introduction

Vaccines containing mRNA encoding SARS-CoV-2 spike antigen, such as Pfizer BNT162b2, can effectively protect people against COVID-19^1-6^. Innate immune sensing of BNT162b2 mRNA by cytosolic RNA sensors immediately after vaccination is required for subsequent activation of spike-specific T-cell and antibody responses^7^. A second dose of BNT162b is sufficient to induce detectable spike-specific antibody and T-cell responses in most individuals, but levels of adaptive immune responses vary widely among individuals^8,9^. Although inter-individual variation in BNT162b2-induced adaptive immunity is associated with several parameters, such as SARS-CoV-2 infection history, age, sex, and ethnicity^9-11^, the cause of this variation remains largely unknown, Recent studies focused on systems biological understanding of human vaccine responses provide important insight into factors associated with inter-individual variation in vaccine-induced adaptive immunity^12-14^. Immune states represented by the composition of immune cells and gene expression profiles in individuals are highly variable, plausibly due to genetic diversity and environmental factors such as gut microbial flora^15-17^. Through comprehensive analysis of immune states of blood cells at baseline and early vaccine responses, specific immune cell populations and transcripts have been identified as correlates of antibody or T-cell responses induced by vaccination against influenza virus, hepatitis B virus, and malaria^18-22^. Moreover, other studies reveal that gut microbiota is also associated with vaccine-induced adaptive immunity^23-25^. Importantly, these factors can be predictors of vaccine responses and may be potential therapeutic targets to improve vaccine responses^26,27^. However, the variability of immune states and gut microbes that is associated with COVID-19 mRNA vaccine responses remains unclear. In this study, using a systems biology approach, we demonstrate that BNT162b2-induced human adaptive immune responses are associated with specific immune and gut microbial parameters.

## Results

### Study design

In this study, we used a systems biology approach based on multi-omics analyses of human blood and stool samples. 96 healthy subjects participated in this study (Supplementary Fig. 1), and data from 95 participants who received two doses of BNT162b2 at a three- to four-week interval were analyzed (data from one participant who was not able to receive the second dose in a timely manner due to severe side effects from the first dose were excluded from the analysis). We collected participant blood samples at five time points (T1-T5) before and after administration of BNT162b2 (Fig. 1). To evaluate the level of vaccine-induced adaptive immunity, we measured the SARS-CoV-2 spike-specific antibody response in plasma and the T-cell response in peripheral blood mononuclear cells (PBMCs). We further used PBMCs for construction of profiles of immune cell populations (cytometry by time of flight (CyTOF) analysis) and mRNA expression (bulk RNA-seq analysis). In addition, to analyze the gut microbiome, we collected stool samples from all subjects once during the participation period (Fig. 1). Through these analyses, we sought to identify immune cell populations, transcripts, and commensal microbial taxa and functions associated with vaccine-induced antibody and T-cell responses.

**Fig. 1.**
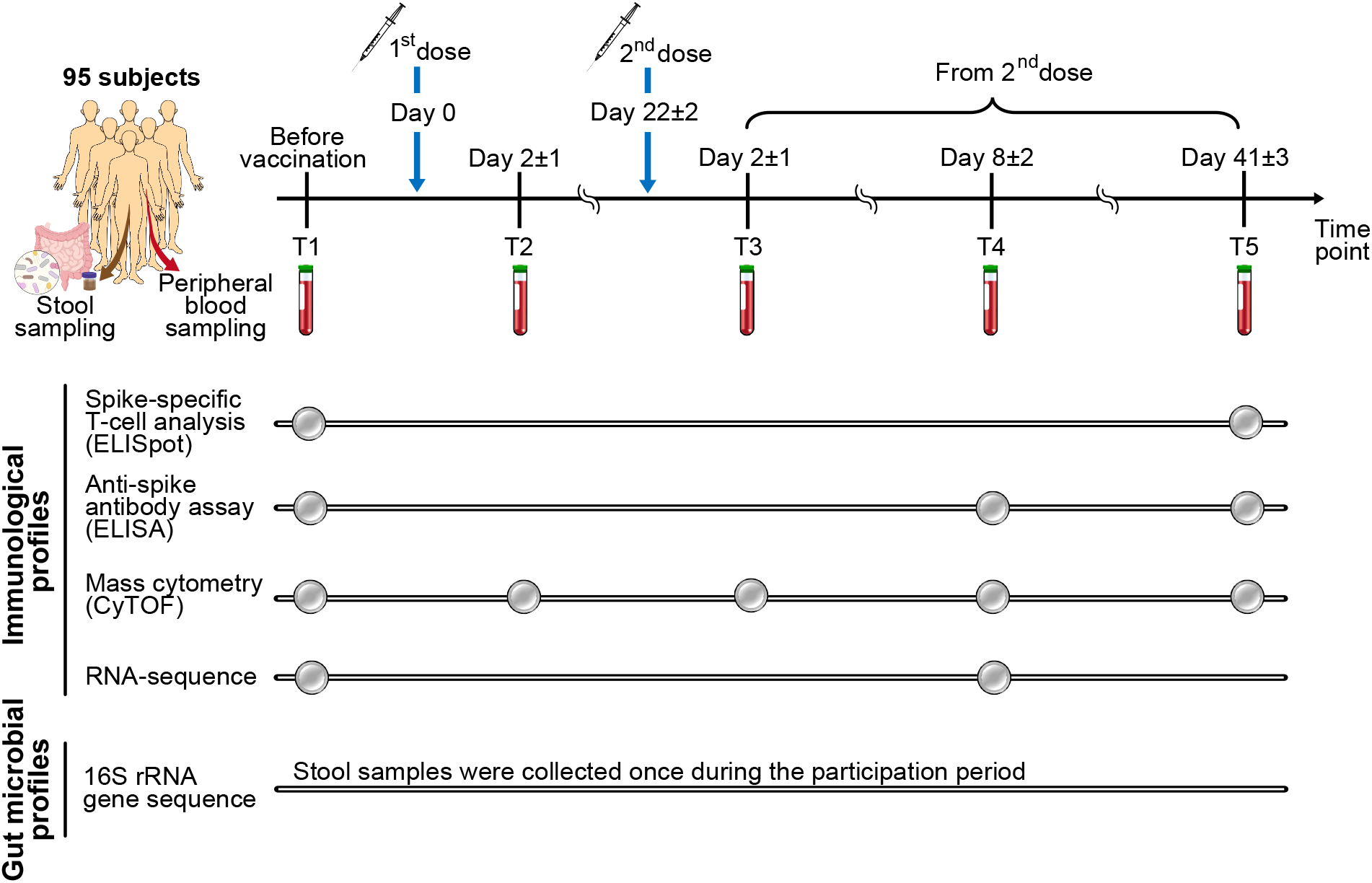
Study design. Schematic diagram showing blood and stool sample collection and analysis performed in this study. Samples from 95 subjects who received two doses of BNT162b2 at 3–4-week intervals were analyzed.

### Inter-individual variation in vaccine-induced adaptive immunity

We first evaluated inter-individual variation in vaccine-induced adaptive immunity by measuring SARS-CoV-2 spike-specific immunoglobulin G (IgG) antibody in plasma and interferon (IFN)-γ-producing T cells in PBMCs by enzyme-linked immunosorbent assay (ELISA) and enzyme-linked immunospot (ELISpot) assay, respectively. We detected an increase in spike-specific antibody and T-cell responses on Day 41±3 after the second dose (T5) in all subjects, but there were significant inter-individual differences in response magnitude (Fig. 2a, b). Subjects who were seropositive for SARS-CoV-2 at baseline (T1) tended to show higher antibody and T-cell responses induced by vaccination. To remove the effect of immunological memory induced by SARS-CoV-2 infection on vaccine-induced adaptive immunity, in subsequent analyses we focused on 86 subjects who were seronegative for SARS-CoV-2 at baseline. Consistent with previous reports, we observed gender-associated differences in antibody and T-cell responses (Fig. 2c, d) and an age-related decline of vaccine-induced antibody responses, but not T-cell responses (Fig. 2e, f). There was no detectable correlation between vaccine-induced antibody and T-cell responses (Fig. 2g). We also measured T-cell responses against four human common cold coronaviruses (HCoV-OC43, 229E, NL63, and HKU1) and found that BNT162b2 vaccination increased the frequency of T cells specific for these HCoVs (Supplementary Fig. 2). Furthermore, there was a significant correlation between T-cell responses against SARS-CoV-2 and these HCoVs (Fig. 2h), indicating that BNT162b2 can induce cross-reactive T cells to HCoVs. Taken together, these results indicate that there are significant inter-individual differences in antibody and T-cell responses elicited by vaccination with BNT162b2.

**Fig. 2.**
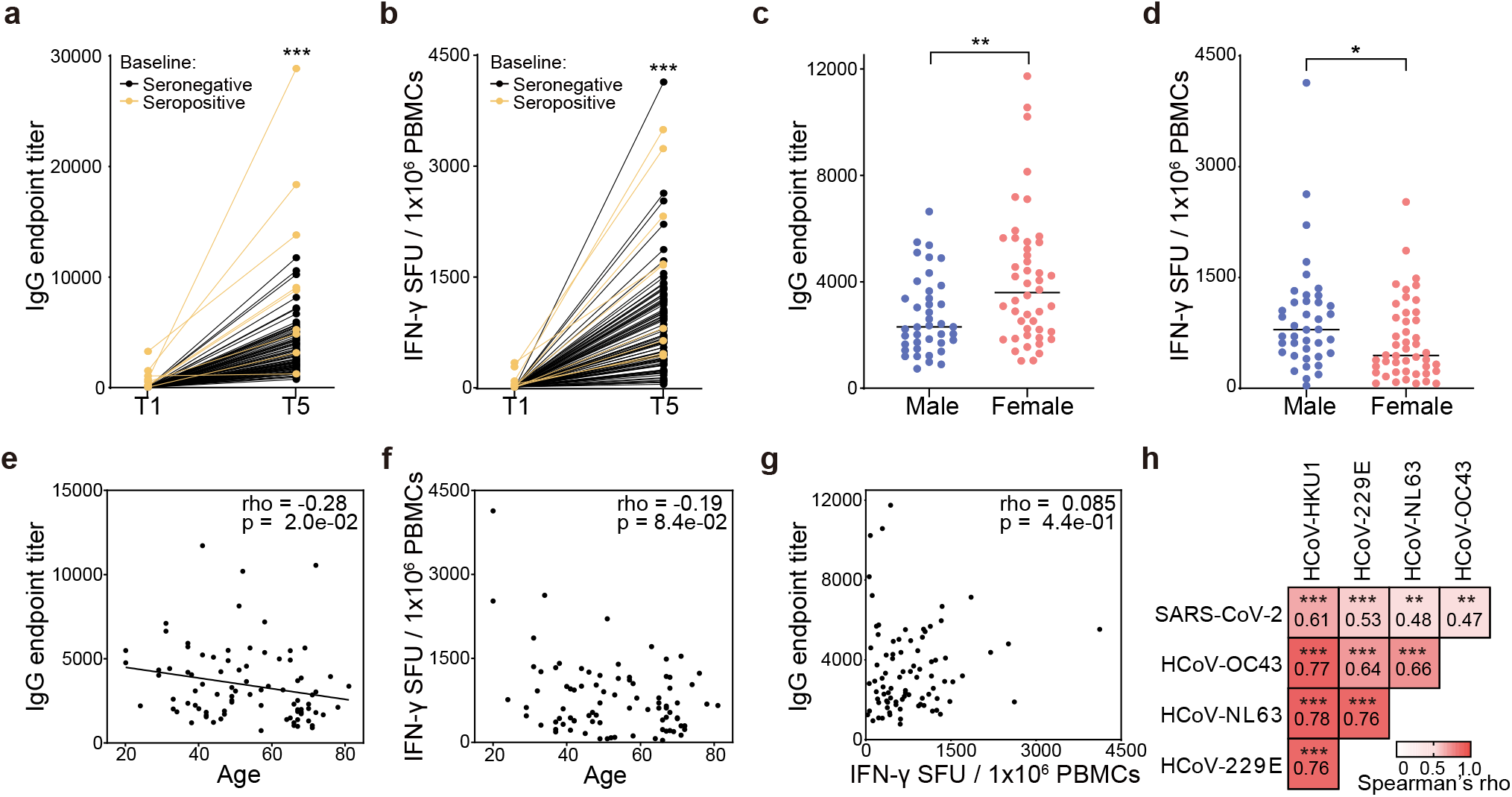
Inter-individual variations in BNT162b2-induced adaptive immunity in our cohort. (**a**)Anti-SARS-CoV-2 spike IgG endpoint titers in plasma at T1 and T5 were measured using ELISA (n = 95). **(b)** IFN-γ-secreting T cells specific for SARS-CoV-2 spike in PBMCs at T1 and T5 were measured with ELISpot assays (n = 95). SFU, spot-forming unit. **(c, d)** BNT162b2-induced antibody responses (anti-SARS-CoV-2 spike IgG endpoint titers at T5) **(c)** and T-cell responses (IFN-γ-secreting T cells specific for SARS-CoV-2 spike in PBMCs at T5) **(d)** in male (n = 40) and female (n = 46) subjects. **(a-d)** p values were calculated by Wilcoxon rank-sum tests (* p < 0.05, ** p < 0.01, *** p < 0.001). **(e, f)** Correlation analysis between age and BNT162b2-induced antibody responses **(e)** or T-cell responses **(f). (g)** Correlation analysis between BNT162b2-induced antibody responses and T-cell responses. **(h)** Heat map showing correlations between vaccine-induced T cell responses against SARS-CoV-2 and HCoVs. **(e-h)** Correlations were analyzed by Spearman’s correlation. p values were corrected with Benjamini–Hochberg FDR correction for multiple tests. Spearman’s rho coefficient and p values are indicated in the plots or in the heat map cells (** p < 0.01, *** p < 0.001).

### Immune cell populations associated with BNT162b2-induced adaptive immunity

To identify cell populations associated with BNT162b2-induced adaptive immune responses, we next performed CyTOF analysis of PBMCs collected at baseline (T1) and after vaccination (T2-T5). Unsupervised dimension reduction and clustering using t-distributed stochastic neighbor embedding (t-SNE) separated PMBCs into 16 major clusters corresponding to subsets of T cells, B cells, natural killer (NK) cells, and monocytes (Fig. 3a). We then compared the frequency of immune cell populations in high- vs low-antibody responders (top 20 vs bottom 20 subjects in antibody titers at T5 among 86 baseline seronegative subjects). This analysis revealed that there were significant differences in the frequency of naïve CD8^+^ T cells and memory CD4^+^ T cells in high- vs low-antibody responders (Fig. 3b). There was a significant positive correlation between the frequency of these cells and vaccine-induced antibody responses (Supplementary fig. 3a), but data adjusted for age and sex did not show such correlations (Fig. 3b). Consistent with previous reports^28-30^, we observed an age-related decline of naïve CD8^+^ T cells (Supplementary Fig. 3b), confirming that aging is the confounding factor affecting both the frequency of naïve CD8^+^ T cells and antibody responses.

**Fig. 3.**
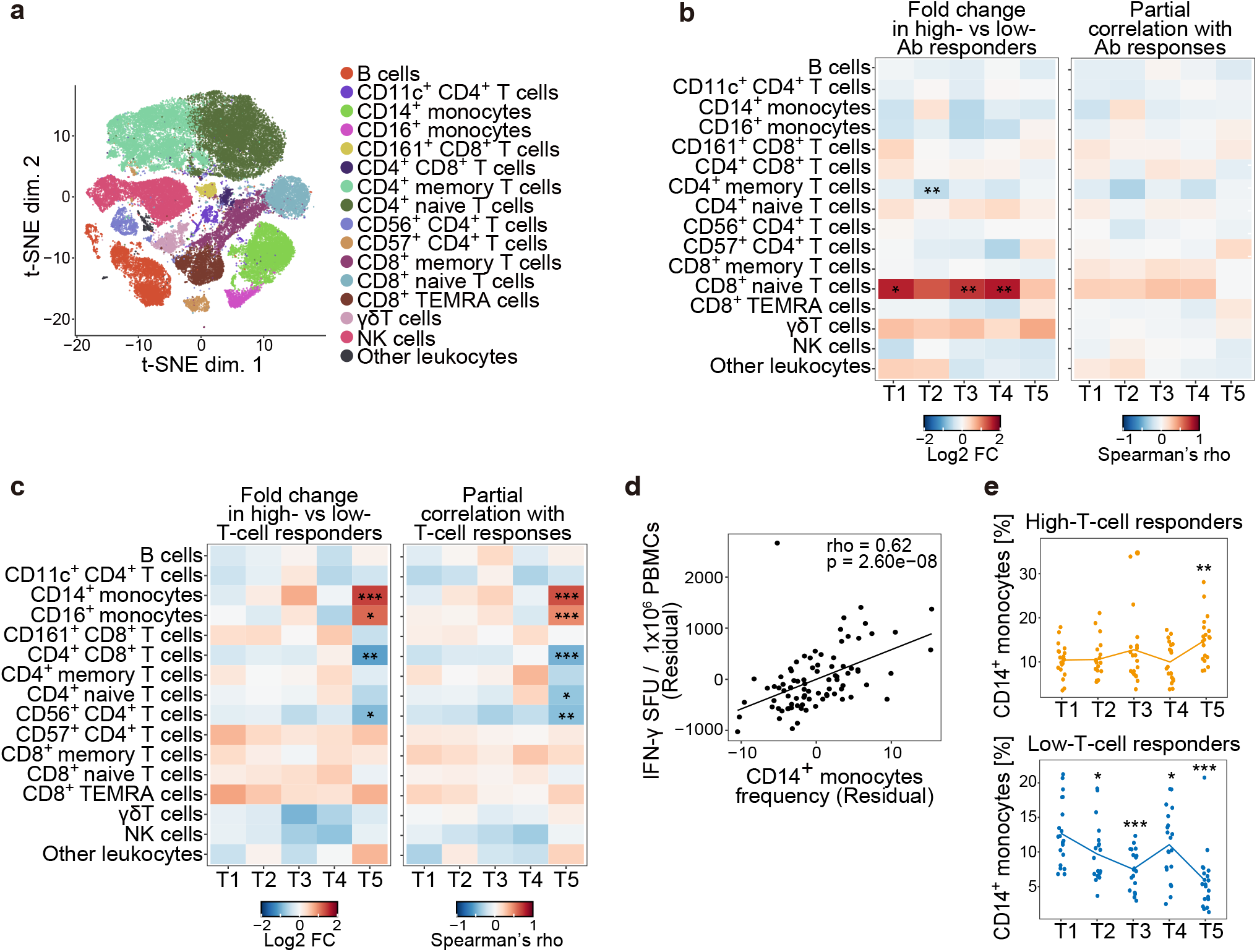
Immune cell populations associated with BNT162b2-induced adaptive immunity. Immune cell populations in PBMCs isolated from baseline seronegative subjects (n = 86) at time points T1-T5 were analyzed with CyTOF. **(a)** t-SNE visualization of CyTOF data of PBMCs at T1 (n = 86). Types of immune cell populations were annotated based on expression of their marker proteins. TEMRA, terminally differentiated effector memory. NK, natural killer. **(b, c)** Left heat map showing differences in the frequency of immune cell populations in high- vs low-antibody (Ab) responders **(b)** or in high- vs low-T-cell responders **(c)** (n = 20 each). Right heat map showing the correlation between the frequency of immune cell populations and vaccine-induced antibody responses **(b)** or T-cell responses **(c)** (n = 86). p values in left panels were calculated by Wilcoxon rank-sum tests with Benjamini–Hochberg FDR correction (* p < 0.05, ** p < 0.01, *** p < 0.001). (d) Scatter plot showing a correlation between the frequency of CD14^+^ monocytes at T5 and vaccine-induced T-cell responses. **(b-d)** Partial correlation analyses with adjustments for age and sex were performed with Spearman’s correlation tests with Benjamini–Hochberg FDR correction (* p < 0.05, ** p < 0.01, *** p < 0.001). **(e)** Kinetics of the frequency of CD14^+^ monocytes in PBMCs during vaccine response. High-T-cell responders (upper panel, n = 20) and low-T-cell responders (lower panel, n = 20) were analyzed. p values were calculated with Wilcoxon signed rank tests with Benjamini–Hochberg FDR correction (** p < 0.01, *** p < 0.0001).

A comparative analysis of frequencies of immune cell populations in high- vs low-T-cell responders (top 20 vs bottom 20 subjects in T-cell responses at T5 among 86 baseline seronegative subjects) showed that the frequency of monocytes was higher in high-T-cell responders than in low responders, while the frequency of several T cell subsets showed the opposite trend, at T5 (Day 41±3 after the second dose) (Fig. 3c). There was a significant correlation between these cell populations and T-cell responses at T5 in the analysis with adjustments for age and sex (Fig. 3c, d). In time course analysis, we observed vaccine-induced increase and decrease in the frequency of monocytes in high-T-cell responders (only at T5) and in low-T-cell responders (from T2 to T5), respectively (Fig. 3e and Supplementary Fig. 3c). Thus, the frequency of monocytes, which changes in the vaccine response, is a positive correlate of vaccine-induced T-cell responses

### Transcripts associated with BNT162b2-induced adaptive immunity

To construct gene expression profiles of PBMCs at baseline and during vaccine response, we performed bulk RNA-seq analysis of PBMCs at T1 (baseline) and T4 (Day 8±2 after the second dose). Of the 86 baseline seronegative subjects, sequence data from 80 (at T1) and 78 (at T4) subjects passed quality control. This analysis revealed that vaccination altered expression of 2296 genes at T4 (Supplementary Fig. 4a). Gene set enrichment analysis (GSEA) revealed that a blood transcription module (BTM) related to plasma cells and B cells was upregulated after vaccination (Supplementary Fig. 4b).

To identify biological pathways associated with BNT162b2-induced adaptive immunity, we next performed GSEA on a ranked gene list based on the correlation with vaccine-induced antibody or T-cell responses. This revealed that a BTM related to the activator protein 1 (AP-1) transcription network was positively and negatively associated with antibody responses (Fig. 4a) and T-cell responses (Fig. 4b), respectively. Furthermore, a comparison between high and low responders in vaccine-induced antibody and T-cell responses showed that 1 gene (at T4) and 130 genes (53 genes at T1 and 77 genes at T4) were differentially expressed (log2 FC > 0.5, adjusted p < 0.05) in high- vs low-antibody responders (Supplementary Fig. 4c) and in high- vs low-T-cell responders (Fig. 4c), respectively. Notably, consistent with the GSEA result, AP-1 transcription factors, such as *FOS, FOSB*, and *JUN* were highly expressed in low-T-cell responders (Fig. 4c).

**Fig. 4.**
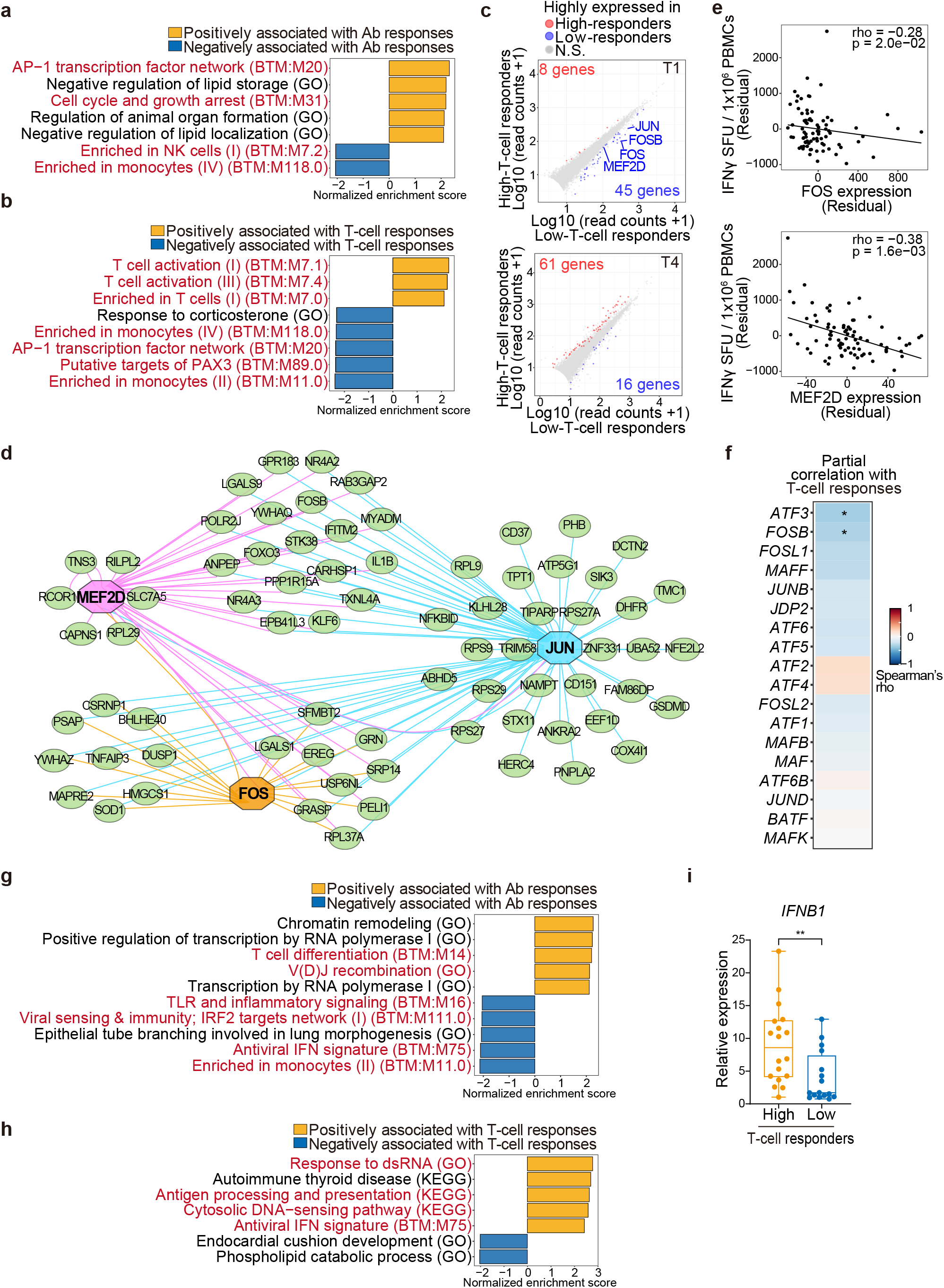
Transcripts associated with BNT162b2-induced adaptive immunity. **(a-e)** Transcriptomes of PBMCs isolated from baseline seronegative subjects (n = 86) at time points T1 and T4 were analyzed by bulk RNA-seq. **(a, b)** GSEA on a ranked gene list based on the spearman’s correlation coefficient between RNA expression and vaccine-induced antibody (Ab) responses **(a)** or T-cell responses **(b)** (n = 86). Immune-related BTM, GO, and KEGG pathways are shown in red. **(c)** Scatter plots showing DEGs (log2 FC > 0.5, adjusted p < 0.05) between high- (n = 18 at T1, n = 19 at T4) and low- (n = 19 at T1, n = 18 at T4) T-cell responders. Red and blue dots indicate DEGs that were highly expressed in high- and low-T-cell responders, respectively. N.S., not significant. **(d)** Gene regulatory network analysis of DEGs between high- and low-T-cell responders. **(e)** Scatter plots showing correlations between vaccine-induced T-cell responses and expression of *FOS* and *MEF2D*. **(f)** Heat map showing correlations between vaccine-induced T-cell responses and expression of AP-1 genes (n = 86). **(e, f)** Partial correlation analyses with adjustments for age and sex were performed with Spearman’s correlation tests with Benjamini–Hochberg FDR correction (* p < 0.05). **(g-i)** PBMCs isolated from baseline seronegative subjects (n = 86) were stimulated with BTN162b2 mRNA for 6 h and analyzed by RNA-seq followed by GSEA on a ranked gene list based on the spearman’s correlation coefficient between RNA expression and vaccine-induced antibody responses **(g)** or T-cell responses **(h)**. Immune-related BTM, GO, and KEGG pathways are shown in red. **(i)** *IFNB1* mRNA expression in high- (n=18) and low- (n=16) T-cell responders was analyzed by qPCR. The p value was calculated with Wilcoxon signed rank test (** p < 0.01).

Gene regulatory network analysis of differentially expressed genes (DEGs) between high- and low-T-cell responders identified *FOS, JUN*, and *MEF2D*, which were highly expressed in low-T-cell responders, as potential regulators for many DEGs (Fig. 4d). Baseline expression of *FOS* and *MEF2D*, but not *JUN*, was inversely correlated with the vaccine-induced T-cell responses in the analysis with adjustments for age and sex (Fig. 4e). Given the correlation between FOS and T cell responses, we assessed whether this is the case for other AP-1 family genes and found that expression of *ATF3* and *FOSB* was inversely correlated with T-cell responses (Fig. 4f). Thus, we identified baseline expression of a subset of AP-1 genes *FOS, FOSB*, and *ATF3* as negative correlates of vaccine-induced T-cell responses.

Next, we sought to investigate whether transcriptomic signatures related to innate immune responses are associated with BNT162b2-induced adaptive immunity. To this end, we performed bulk RNA-seq analysis of PBMCs stimulated with BNT162b2 mRNA for 6 h *ex vivo*, because relatively large time lags in our blood sampling did not allow us to evaluate dynamic gene expression in BNT162b2-induced innate immunity. BNT162b2 mRNA stimulation upregulated genes related to type I interferon (IFN) responses (Supplementary Fig. 4d, e). GSEA revealed that a BTM related to type I IFN responses was negatively and positively associated with antibody responses (Fig. 4g) and T-cell responses (Fig. 4h), respectively. Consistent with this, qPCR analysis showed that *IFNB1* expression was significantly higher in high-T-cell responders than low responders (Fig. 4i). These data suggest that expression of type I IFN genes in the early innate immune response is positively associated with BNT162b2-induced T-cell responses.

### Baseline FOS expression is negatively associated with early T-cell responses to BNT162b2 mRNA

To investigate whether and how baseline expression of AP-1 transcription factors is associated with early vaccine response, we performed single-cell RNA-seq (scRNA-seq) analysis of PBMCs of subjects who exhibited high or low *FOS* expression in the bulk RNA-seq analysis (high- and low-*FOS* subjects, n=4 each) in the absence or presence of *ex vivo* stimulation with BNT162b2 mRNA. This experimental setting allowed us to evaluate the association between *FOS* and other genes expression at baseline and in early innate immune response (6 and 16 h after BNT162b2 mRNA stimulation) in specific cell populations (Fig. 5a). Unsupervised clustering identified 9 major immune cell populations whose frequencies were comparable between high- and low-*FOS* subjects (Fig. 5b). BNT162b2 mRNA stimulation upregulated genes related to RIG-I-like receptor signaling and type-I IFN response, particularly in the monocyte population (Fig. 5c and Supplementary Fig. 5a).

**Fig. 5.**
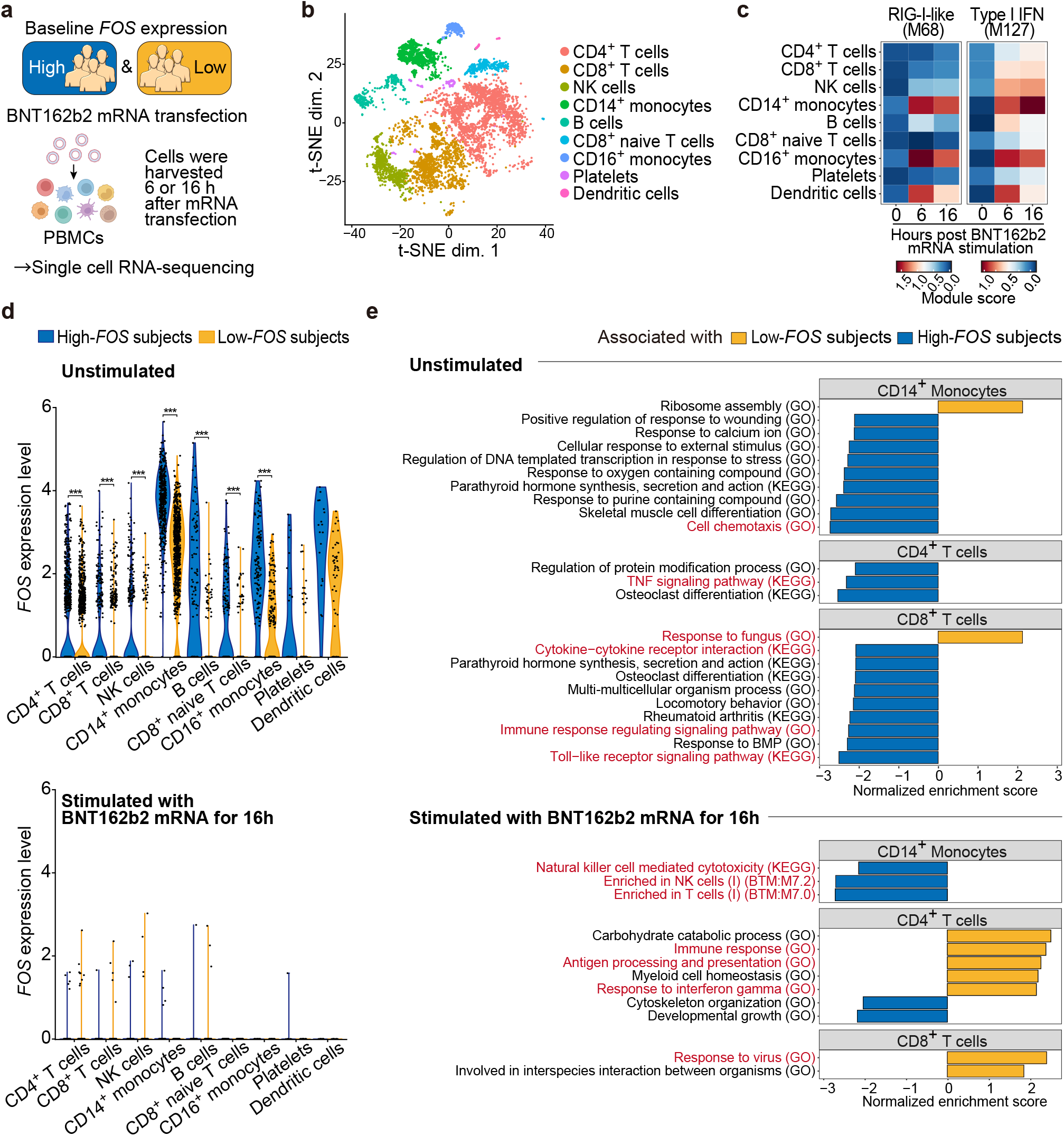
Baseline *FOS* expression is negatively associated with early T-cell responses to BNT162b2 mRNA. PBMCs isolated from subjects who exhibited high or low *FOS* expression in the bulk RNA-seq analysis (high and low *FOS* subjects, n = 4 each) were unstimulated or stimulated with BNT162b2 mRNA for 6 or 16 h, followed by scRNA-seq analysis. **(a)** Schematic illustrating the experimental design of scRNA-seq of high- and low-*FOS* subjects. **(b)** t-SNE visualization of scRNA-seq data of unstimulated PBMCs. Data from all eight subjects (high and low *FOS* subjects, n = 4 each) were pooled and visualized. **(c)** Module score analysis of genes differentially expressed between immune cell populations stimulated with BNT162b2 mRNA and unstimulated populations. **(d)** Violin plots showing expression of *FOS* in PBMCs unstimulated (upper panel) and stimulated with BNT162b2 mRNA for 16 h (lower panel). *FOS* expression levels in each immune cell population were compared between high- and low-*FOS* subjects (n = 4 each). p values were calculated with Wilcoxon rank-sum tests with Benjamini–Hochberg FDR correction (*** p < 0.001). **(e)** GSEA on a ranked gene list based on the fold change in expression in CD14^+^ monocytes, CD4^+^ T cells, and CD8^+^ T cells unstimulated or stimulated with BNT162b2 mRNA for 16 h between high- and low-*FOS* subjects. Immune-related BTM, GO, and KEGG pathways are shown in red.

We found that *FOS* was expressed all over the immune cell populations that we detected in unsupervised clustering analysis, with the highest expression in CD14^+^ monocytes, in the absence of BNT162b2 mRNA stimulation (Fig. 5d). As expected, *FOS* expression was significantly higher in high-*FOS* subjects than low-*FOS* subjects (Fig. 5d). However, *FOS* expression was significantly reduced in response to BNT162b2 mRNA stimulation in most PBMC subpopulations (Fig. 5d and Supplementary Fig. 5b). To investigate genes associated with baseline *FOS* expression in each cluster, we next performed GSEA on a ranked gene list based on fold changes in expression between high- and low-*FOS* subjects. This showed that GO terms related to baseline immunity, such as chemotaxis in CD14^+^ monocytes, the tumor necrosis factor (TNF) signaling pathway in CD4^+^ T cells, and the Toll-like receptor signaling pathway in CD8^+^ T cells, were associated with high-*FOS* subjects at baseline (Fig. 5e and Supplementary Fig. 5c). In contrast, upon BNT162b2 mRNA stimulation, GO terms related to T cell activation, such as response to IFN-γ in CD4^+^ T cells and responses to virus in CD8^+^ T cells, were associated with low-*FOS* subjects (Fig. 5e and Supplementary Fig. 5c). Taken together, these results indicate that *FOS* expression is positively associated with expression of genes related to baseline immune cell activity, but it is negatively associated with that related to T cell activation upon BNT162b2 mRNA stimulation.

### Gut microbes associated with BNT162b2-induced adaptive immunity

To assess the association between commensal gut microbes and vaccine-induced adaptive immunity, we next performed 16S ribosomal RNA gene sequencing analysis using stool samples of subjects.

There was no difference in Shannon’s diversity index in high- vs low-antibody responders and in high- vs low-T-cell responders (Supplementary Fig. 6a). Linear discriminant analysis effect size (LEfSe) analysis identified 23 taxa and 11 taxa that were differentially enriched in high- vs low-antibody responders and in high- vs low-T-cell responders, respectively (Fig. 6a, b). However, there were no significant correlations between these taxa and vaccine-induced antibody or T cell responses in analysis with adjustments for age, sex, and stool sampling timing (Supplementary Fig. 6b, c).

**Fig. 6.**
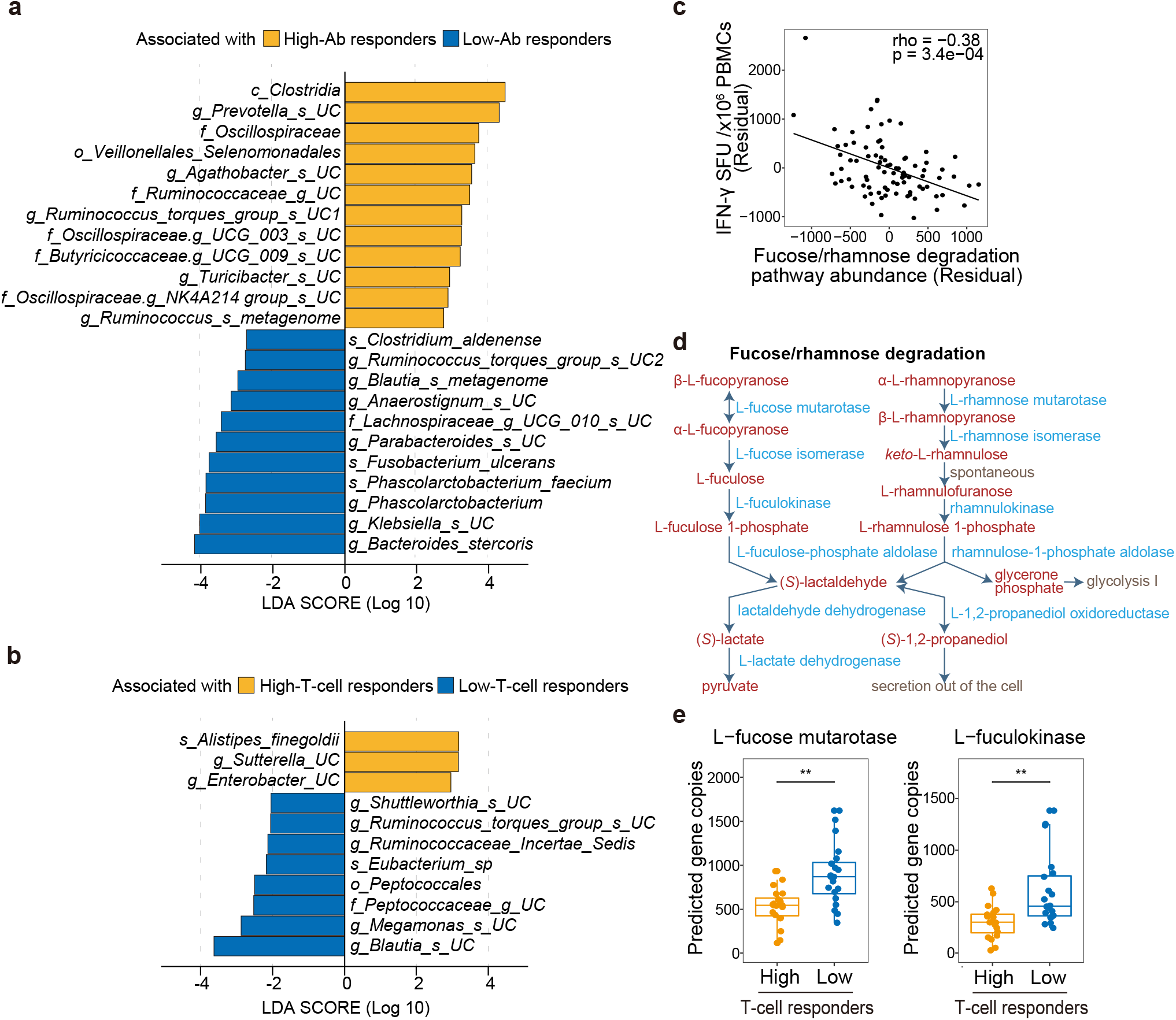
Gut microbes associated with BNT162b2-induced adaptive immunity. Microbiomes of stool samples were analyzed by 16S ribosomal RNA gene sequencing (n = 86). **(a, b)** LEfSe analysis of gut microbes that were differentially abundant in high- vs low-antibody (Ab) responders **(a)** and in high- vs low-T-cell responders **(b)** (n = 20 each). o, order; f, family; g, genus; s, species; UC, unclassified. **(c)** Scatter plot showing a correlation between the gut microbial fucose/rhamnose degradation pathway and vaccine-induced T-cell responses (n = 86). Partial correlation analysis with adjustments for age, sex, and stool sampling timing was performed with Spearman’s correlation tests. **(d)** Schematic showing the fucose/rhamnose degradation pathway. Metabolites and enzymes involved in the pathway are shown in red and blue, respectively. **(e)** Analysis of the abundance of predicted gene copies for *L-fucose mutarotase* and *L-fuculokinase* in high- and low-T-cell responders (n = 20 each). p values were calculated with the Wilcoxon rank-sum tests with Benjamini–Hochberg FDR correction (** p < 0.01).

We next searched for functions of gut microbiota that are associated with vaccine-induced adaptive immunity using a metagenome prediction tool, phylogenetic investigation of communities by reconstruction of unobserved states (PICRUSt2). This analysis revealed that the fucose/rhamnose degradation pathway of gut microbiota was inversely correlated with vaccine-induced T-cell responses (Supplementary Fig. 6d). Partial correlation analysis confirmed that the correlation between the fucose/rhamnose degradation pathway and T-cell responses was independent of age, sex, and fecal sampling timing (Fig. 6c). The fucose/rhamnose degradation pathway converts fucose to lactaldehyde, which in turn is converted to (*S*)-1,2-propanediol or pyruvate (Fig. 6d). Among enzymes involved in this pathway, abundances of genes encoding *L-fucose mutarotase* and *L-fuculokinase* were significantly higher in microbiomes of low-T-cell responders (Fig. 6e and Supplementary Fig. 6e). Furthermore, we found that *Blautia*, which was enriched in low-T-cell responders (Fig. 6b), was a dominant taxon that encodes *L-fucose mutarotase* (Supplementary Fig. 6f, g). Taken together, these data indicate that the gut microbial fucose/rhamnose degradation pathway is a negative correlate of vaccine-induced T-cell responses.

### The gut microbial fucose/rhamnose degradation pathway is associated with AP-1 expression

Finally, we investigated whether the gut microbial fucose/rhamnose degradation pathway is associated with baseline expression of transcription factors that we identified as correlates of vaccine-induced T-cell responses. This showed that the gut microbial fucose/rhamnose degradation pathway was positively correlated with baseline *FOS, FOSB*, and *ATF3* expression in PBMCs (Fig. 7a-d). The fucose/rhamnose degradation pathway generates (*S*)-1,2-propanediol and pyruvate, which in turn leads to generation of short-chain fatty acids (SCFAs) (Fig. 7e). SCFAs derived from intestinal bacteria contribute to modulating host immune responses by inducing colonic regulatory T cell differentiation^31-33^. Furthermore, SCFAs induce production of prostaglandin E2 (PGE2), which upregulates AP-1 expression^34^. Therefore, we assessed whether SCFAs promote PGE2 expression in PBMCs. This showed that SCFAs, but not (*S*)-1,2-propanediol, significantly increased expression of *COX2* (Fig. 7f), which encodes an enzyme catalyzing production of prostaglandins. Furthermore, prostaglandin E2 (PGE2) treatment enhanced expression of *FOS* in PBMCs (Fig. 7g). These results suggest a potential functional link from the gut microbial fucose/rhamnose degradation pathway to AP-1 gene expression in PBMCs.

**Fig. 7.**
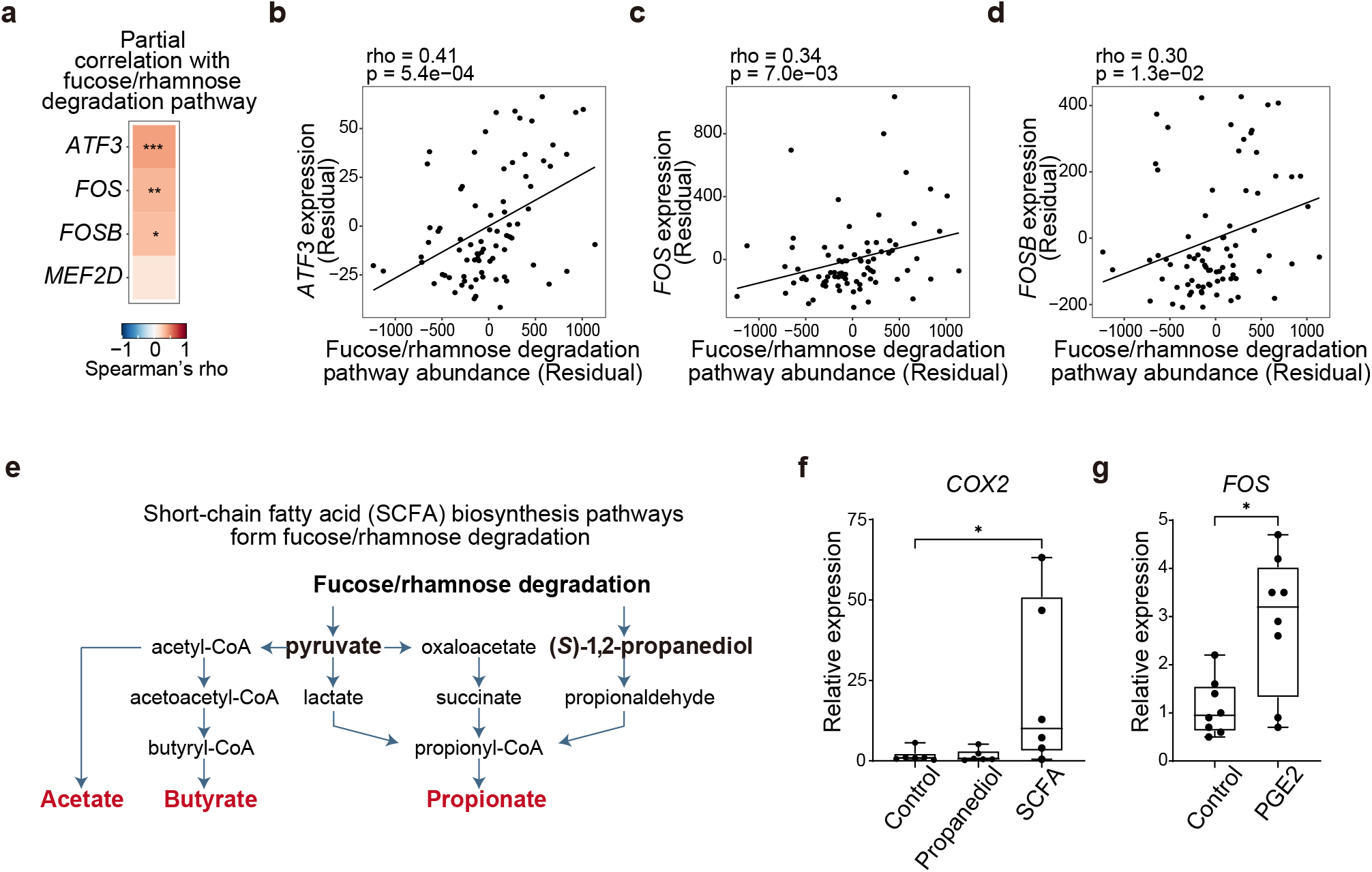
The gut microbial fucose/rhamnose degradation pathway is associated with AP-1 expression. **(a)** Heat map showing the correlation between the gut microbial fucose/rhamnose degradation pathway and transcription factors associated with BNT162b2-induced T-cell responses (n = 86). **(b-d)** Scatter plots showing the correlation between the gut microbial fucose/rhamnose degradation pathway and baseline expression of *ATF3* **(b)**, *FOS* **(c)**, and *FOSB* **(d)** (n = 86). **(a-d)** Partial correlation analyses with adjustments for age, sex, and stool sampling timing were performed with Spearman’s correlation tests with Benjamini–Hochberg FDR correction (* p < 0.05, ** p < 0.01, *** p < 0.001). **(e)** Schematic showing the production of SCFAs from the fucose/rhamnose degradation pathway. SCFAs are shown in red. **(f)** qPCR analysis of *COX2* mRNA levels in PBMCs untreated or treated with (*S*)-1,2-propanediol or SCFAs for 18 h (n = 6). p values were calculated using the Friedman test followed by Dunn’s multiple comparison test (* p < 0.05). **(g)** qPCR analysis of *FOS* mRNA levels in PBMCs untreated or treated with PGE2 for 18 h (n = 8). The p value was calculated with the Wilcoxon signed rank test (* p < 0.05).

## Discussion

In this study, we identified various human immune cell populations and transcripts as well as gut bacterial taxa and functional pathways that are associated with BNT162b2-induced vaccine responses using a systems biology approach. Notably, the baseline transcription module related to the AP-1 transcription factor network was positively associated with BNT162b2-induced antibody response and negatively associated with T-cell responses. Consistent with this, the T cell response was inversely correlated with baseline expression of AP-1 genes (*FOS, FOSB*, and *ATF3*). Furthermore, the gut microbial fucose/rhamnose pathway was inversely correlated with T-cell responses. These findings advance our understanding of the contribution of immune and microbial factors to inter-individual variations in vaccine-induced adaptive immunity.

This study provides new insight into the role of AP-1 genes in vaccine-induced T-cell responses. We observed that AP-1 expression in PBMCs rapidly decreased upon *ex vivo* stimulation with BNT162b2 mRNA, which is consistent with a recent report that expression of AP-1 genes such as *FOS* and *ATF3* was diminished in CD14^+^ monocytes by BNT162b2 vaccination^35^. Interestingly, the AS3-adjuvanted H5N1 pre-pandemic influenza vaccine also induces a decrease of AP-1 genes expression in monocytes through epigenetic silencing, which likely inhibits AP-1-regulated cytokine expression^36^. However, how the difference in baseline AP-1 expression affects vaccine response remains unknown. We found that *FOS* expression, which is inversely correlated with vaccine-induced T-cell responses, is positively associated with transcription modules related to baseline activity of CD14^+^ monocytes and T cells. Furthermore, baseline FOS expression is negatively associated with transcription modules related to T cell activation upon BNT162b2 mRNA stimulation *ex vivo*. These data suggest that baseline expression of FOS and other AP-1 factors in T cells and/or FOS-dependent control of baseline immune cell activity may inhibit T-cell activation mediated by mRNA vaccines.

Our results suggest a novel functional link between the gut microbial fucose/rhamnose degradation pathway and the host immune system. The fucose/rhamnose degradation pathway can promote generation of SCFAs by several mechanisms, including cross-feeding of (*S*)-1,2-propanediol, a metabolic end-product of this pathway, between gut commensal bacteria resulting in production of propionate^37^. SCFAs have immunomodulatory functions, such as promoting mucosal Treg generation^31-33^. Given these, our finding suggests that fucose/rhamnose degradation may result in an increase of SCFAs, which in turn facilitates Treg generation, thereby inhibiting vaccine-induced T-cell responses. Furthermore, our data suggest that SCFAs upregulate PGE2 production through upregulation of *COX2* expression, which in turn upregulates *FOS* expression in PBMCs. Future studies will need to further explore the clinical significance and molecular mechanisms of interactions between the fucose/rhamnose pathway and vaccine-induced T-cell responses.

Our CyTOF analysis revealed a significant difference in the frequency of monocytes on Day 41 after the second dose between low- and high-T-cell responders. We observed a decrease of monocytes for at least two months after BNT162b2 vaccination in low-T-cell responders, but not in high responders. Conversely, there was an increase of monocytes between Days 8 and 41 after the second dose only in high-T-cell responders. These observations indicate remarkable heterogeneity in monocyte response induced by BNT162b2 vaccination. Infection and vaccination can affect monocyte development, homeostasis, and migration, thereby altering the frequency of monocytes in the blood^38,39^. Interestingly, vaccination with BCG, AS3-adjuvanted H5N1 pre-pandemic influenza (H5N1+AS03) vaccine, or HIV vaccine induces innate memory monocytes that provide protection against non-related^36,40^ and related viruses^41^. Epigenetic changes induced by H5N1+AS03 are maintained in monocytes for at least 6 months, suggesting a long-lasting trained immunity^36^. Accordingly, it would be interesting to assess whether BNT162b2-induced changes in monocyte frequency are associated with memory monocyte generation and whether this affects host defense.

This study successfully identified multiple correlates of BNT162b2-induced adaptive immunity, but several shortcomings in the sampling scheme and experimental design may have prevented identification of other correlates. First, the relatively small sample size and the ethnic and geographic bias of participants in this study may have limited identification of correlates of adaptive immune responses. This may be one of the reasons why several enterobacterial taxa correlated with BNT162b2-induced antibody responses were identified in another study^25^, but not in our study.

Second, time lags in blood sampling over several days may have impeded identification of correlates of adaptive immunity that change dynamically in short time windows, such as genes induced by innate immunity. However, this issue was partly addressed by our RNA-seq analysis of PBMCs stimulated with BNT162b2 mRNA *ex vivo*. Third, we used the level of IFN-γ-secreting T cells as an indicator of T-cell responses for simple and accurate measurement by ELISpot assay, but analysis of CD4^+^ and CD8^+^ effector T cell subsets may be more informative. Fourth, high-throughput, scRNA-seq analysis and higher resolution of cell phenotyping by CyTOF will be required for a more comprehensive understanding of inter-individual variations of vaccine-induced adaptive immunity.

In summary, we discovered several new immune and microbial parameters at baseline and in the vaccine response that are associated with BNT162b2-induced antibody and T-cell responses, which provide insight into mechanisms of inter-individual variation in adaptive immunity. Our data suggest a key role of baseline AP-1 expression and the gut microbial fucose/rhamnose degradation pathway in inter-individual variation in mRNA vaccine-induced T-cell responses. Future studies should address the potential of these factors as baseline predictors of vaccine outcome and as therapeutic targets to improve vaccine responses.

## Methods

### Subjects

The study was approved by the Okinawa Institute of Science and Technology, Graduate University (OIST) human subjects ethics committee (application HSR-2021–001). Ninety-six Japanese healthy volunteers (42 men and 53 women; average age, 52.4 ± 14.9 years; age range: 20–81 years) were recruited in Okinawa, Japan, between May 2021 and August 2021. All participants provided informed written consent. 25 mL of peripheral blood was collected at each sampling. Stool samples were also collected from all participants once during the participation period.

### PBMCs and plasma collection

Blood samples were collected in heparin-coated tubes (TERUMO; VP-H100K). PBMCs and plasma were separated using Leucosep tubes pre-filled with Ficoll-Paque Plus (Greiner; 163288), as previously described^42^. Briefly, 25 mL of blood and 12 mL of AIM-V medium (Thermo; 12055091) were added to Leucosep tubes and centrifuged at 1,000 g at room temperature for 10 min, and the upper yellowish plasma solution and the white layer containing PBMCs were collected. PBMCs were then washed twice with 22 mL of AIM-V medium with centrifugation at 600 g (for the first wash) or 400 g (for the second wash) for 7 min. PBMC pellets were resuspended in 500 μL of CTL test medium (Cellular Technology Limited (CTL); CTLT-010). Fresh PBMCs were used for IFN-γ ELISpot assays. Plasma was collected and stored at -20°C, and PBMCs were stored in liquid nitrogen until use.

### SARS-CoV-2 antibody ELISA

Anti-SARS-CoV-2 spike IgG ELISA assays were performed as previously described^43,44^ with minor modifications. Briefly, 96-well plates were coated with 2-4 μg/mL HexaPro^45^ spike protein overnight at 4°C. Concentration was adjusted as necessary to optimize positive control signal reproducibility across protein purification batches. After blocking with 200 μL of PBST plus 3% milk, prepared serial dilutions of sera in PBST plus 1% milk were transferred to ELISA plates. Antibody incubation steps were carried out in an incubator at 20 °C. All other steps were carried out as described previously^43^. For data analysis, the background value was set at an OD492 of 0.2 AU, and the endpoint titer was calculated using Prism 7 (GraphPad).

### IFN-γ ELISpot assay

Peptide pools for SARS-CoV-2 S (JPT; PM-WCPV-S-1), HCoV-OC43 (GSC; PR30011), HCoV-NL63 (JER; PM-NL63-S-1), HCoV-229E (GSC; RP30010), and HCoV-HKU1 (JER; PM-HKU1-S-1) proteins dissolved in DMSO (500 μg/mL) were used for cell stimulation. IFN-γ ELISpot assays were performed using Human IFN-γ Single-Color Enzymatic ELISpot kits (CTL; hIFNgp-2 M), according to the manufacturer’s instructions. Briefly, freshly isolated PBMCs (2.5 × 10^5^ cells per well) were stimulated with 1 μg/mL peptide solutions for each SARS-CoV-2 protein for 18 h. For each sample analysis, negative controls (cells treated with equimolar amounts of DMSO) and positive controls (cells treated with 20 ng/mL phorbol 12-myristate 13-acetate (PMA) and 100 ng/mL ionomycin) were included. After incubation, plates were washed and developed with detection reagents included in the kits. Spots were counted using a CTL ImmunoSpot S6 Analyzer. Antigen-specific spot counts were determined by subtracting background spot counts in a negative control well from wells treated with peptide pools.

### CyTOF immunophenotyping

Cryopreserved PBMCs were thawed, centrifuged for 5 min at 440 g, and resuspended in TexMACS Medium (Miltenyi Biotec). Cells were then treated with DNase I (100 U/mL) in the presence of 5 mM MgCl2 for 15 min, centrifuged and resuspended in staining buffer, followed by barcoding with different combinations of Maxpar human anti-CD45 antibodies labeled with 106Cd, 110Cd, 111Cd, 112Cd, 113Cd, or 114Cd. (Fluidigm). 18-20 barcoded PBMC samples were pooled (1 × 10^5^ cells/sample) and immunostained using a Maxpar Direct Immune Profiling Assay kit (Fluidigm) according to the manufacturer’s protocol. PBMC samples were washed three times with Cell Acquisition Solution (CAS) or CAS plus buffer (Fluidigm) and resuspended in the same buffer containing a 1/10 dilution of EQ beads (Fluidigm). Samples were analyzed (an average of 5 × 10^4^ events/sample) with a Helios mass cytometer system (Fluidigm).

### CyTOF data analysis

FCS files were normalized using EQ beads and concatenated. Then the files were de-barcoded using the barcode key file (Key_Cell-ID_20-Plex_Pd.csv) in the Fluidigm acquisition software (v. 6.7.1014). Clean-up gates for live single cells and elimination of non-cell signals were manually conducted using the web-plat software, Cytobank (v.9.1). To correct batch effects across CyTOF runs, signal intensities were normalized using cyCombine^46^. Data were analyzed using a previously described R-based pipeline^47^. In brief, data were imported and transformed for analysis using the read.flowSet function from the flowCore package^48^ and the prepData with option (cofactor = 5) function from the CATALYST (https://github.com/HelenaLC/CATALYST) package, respectively. Clustering was based on the fastPG^49^ algorithm with default parameters. These clusters were visualized using t-SNE and subsequently annotated based on protein markers expression.

### Bulk-RNA seq

Cryopreserved PBMCs were thawed and centrifuged for 5 min at 440 g, and total RNA was isolated using Isospin cell and tissue RNA kit (Nippon Gene) or an RNAdvance v2 kit (Beckman Coulter) according to manufacturer instructions and quantified with an RNA HS Assay Kit (Thermo Fisher) and a Qubit Flex Fluorometer (Thermo Fisher). For transcriptome analysis, 10 ng of RNA were used for library preparation with a QuantSeq 3′ mRNA-Seq Library Prep Kit FWD for Illumina (Lexogen) according to the manufacturer’s protocol for low-input RNA samples. To generate single-nucleotide polymorphism (SNP) calls for several donors whose samples were analyzed by scRNA-seq, cDNA libraries were prepared from 500 ng of RNA using a Collibri Stranded RNA Library prep Kit (Thermo Fisher) according to the manufacturer’s protocol for degraded RNA samples. Libraries were quantified with a Qubit 1x dsDNA HS Assay Kit (Thermo Fisher) and a Qubit Flex Fluorometer (Thermo Fisher), and quality was assessed using D1000 ScreenTape and High Sensitivity D5000 ScreenTape with a Tapestation 2200 (Agilent). Pooled libraries were sequenced on a Novaseq 6000 instrument (Illumina) with 1×100-bp reads for transcritome analysis and 2×150-bp reads for generation of SNP calls at the Sequencing Section at OIST.

### Bulk RNAseq data processing

To evaluate data quality, we applied FastQC (v.0.11.9) (www.bioinformatics.babraham.ac.uk/projects/fastqc/). Reads were further processed to remove adaptor and low-quality sequences using Trimmomatic^50^ (v.0.39) software with the options (SLIDINGWINDOW:4:20 LEADING:20 TRAILING:20 MINLEN:20 HEADCROP:12). To align reads to the GRCh38 reference genome (Homo_sapiens.GRCh38.dna.primary_assembly.fa file downloaded from Ensembl), we used HISAT2^51^ (v.2.2). We counted the number of reads overlapping the genes in GENCODE (v.30) reference transcriptome annotations using featureCounts from Subread^52^ (v.2.0.1) with flags (-s 1 -t gene). The samples with fewer than 300,000 total reads were excluded from the analysis. To detect differentially expressed genes between the high- and low-Ab or T-cell responders, we first filtered transcripts with an average read count of less than 5 and analyzed statistical significance with the Wald test using DESeq2^53^ (v.1.34.0). Gene set enrichment analysis based on blood transcriptional module (BTM), Kyoto Encyclopedia of Genes and Genomes (KEGG), and Gene Ontology collection (GO) was performed using the clusterProfiler^54^ package (v.4.2.2). To predict regulators that explain the observed differential transcriptional program between the two groups, we used iRegulon^55^ (v.1.3) through the Cytoscape (v.3.9.1) visualization tool. Analysis was performed on the putative regulatory region of 20 kb centered around the transcription start site using default settings.

### SNP calling

Sequencing reads were adaptor- and quality-trimmed and then aligned to the human genome using the Hisat2 aligner. SNP calls were generated using a previously published protocol^56^. In brief, we used SAMtools^57^ (v.1.12) to remove duplicates (command markdup). Then, we applied the BEDtools^58^ (v.2.26.0) intersect to identify and remove SNPs in imprinted genes (http://www.geneimprint.org/ accessed: 3 January 2022) and SNPs in repeats (RepeatMasker annotation downloaded from the UCSC Genome Browser). Genotypes were obtained with SAMtools mpileup with options (-A -q 4 -t AD, DP) and BCFtools^59^ (v.1.11-1) call (with options –m --O b -f GQ), using uniquely mapped reads. We used VCFtools^60^ (v.0.1.16-2) to select SNPs with a depth ≥ 10 with options (-minDP 10) and a genotype quality ≥ 20 with options (-minGQ 20).

### *Ex vivo* PBMC stimulation with BNT162b2 mRNA

The BNT162b2 cDNA sequence, including 5’ and 3’ untranslated regions^61^, was synthesized by IDT and cloned into pCDNA3.1 (Thermo Fisher). Using PCR-amplified BNT162b2 cDNA with an upstream T7 promoter as a template, *in vitro* transcription of BNT162b2 mRNA was performed with a HiScribe T7 ARCA mRNA Kit with tailing (NEB) with 2.5 mM N1-Methylpseudouridine-5’-triphosphate nucleoside analog (TriLink BioTechnologies) instead of unmodified UTP. BNT162b2 mRNA was purified using a Monarch RNA cleanup kit (NEB) and dissolved in nuclease-free water.

Cryopreserved PBMCs were thawed, centrifuged for 5 min at 440 g, and resuspended in TexMACS Medium (Miltenyi Biotec). PBMCs were seeded into a 96-well plate (10^6^ cells/well) and stimulated by transfection with BNT162b2 mRNA (200 ng/well) using Lipofectamine MessengerMAX (Thermo Fisher) according to the manufacturer’s instructions. Cells were harvested 6 or 16 h after mRNA transfection.

### scRNA-seq

PBMCs unstimulated or stimulated with BNT162b2 mRNA for 6 h or 16 h were used for analysis. Cells from 8 subjects were pooled in equal numbers and resuspended in ice-cold PBS with 0.04% BSA at a final concentration of 1000 cells/μL. Single-cell suspensions were then loaded on the 10X Genomics Chromium Controller with a loading target of 20,000 cells. Libraries were generated using a Chromium Next GEM Single Cell 5′ v2 (Dual Index) Reagent Kit according to the manufacturer’s instructions. A Quantitative PCR Bio-Rad T100 Thermal Cycler (Biorad) was used for a reverse transcription reaction. All libraries were quality controlled using a Tapestation (Agilent) and quantified using a Qubit Fluorometr (ThermoFisher). Libraries were pooled and sequenced on an Illumina NovaSeq platform (Illumina) using the following sequencing parameters: read1-26-cycle, i7-10, i5-10, read2-90 with a sequencing target of 20,000 reads per cell RNA library.

### scRNA-seq data analysis

The CellRanger Single-Cell Software Suite (10x Genomics) was used to perform barcode processing and transcript counting after alignment to the GRCh38 reference genome with default parameters. To match single cells in the 10x RNAseq data to each donor and identify doublets, we used the software package demuxlet^62^, which uses variable SNPs between pooled individuals. To further analyze scRNAseq data, we used the Seurat^63^ R package. Cells expressing >5% mitochondrial gene counts or expressing less than 500 genes were discarded using the subset function. Then, the NormalizeData and FindVariableFeatures functions were applied to each dataset before FindIntegrationAnchors, IntegrateData and ScaleData were called to combine and scale the data. Unsupervised clustering was applied in each dataset as follows: (i) The top variant genes selected by FindVariableFeatures were used as input for principal components analysis (PCA) to reflect major biological variation in the data. (ii) The top 15 principal components were used for t-SNE dimensional reduction with the RunTSNE function and unsupervised clustering. Specifically, the FindClusters function was used to cluster cells. (iii) After cell clusters were determined, marker genes for each cluster were identified by the FindAllMarkers function with default parameters. The AddModuleScore function was used to calculate the module score in each cell. Plots of expression of specific transcripts were created using the FeaturePlot function. To find differentially expressed genes between high- and low-*FOS* groups, we used the FindMarkers function with the MAST algorithm^64^. Gene set enrichment analysis based on BTM, KEGG, and GO was performed using the clusterProfiler R package.

### 16S rRNA gene sequencing

DNA was extracted from stool samples using QIAmp Fast DNA Stool Mini Kit (Qiagen). 16S rRNA V3 and V4 regions were amplified by PCR using Kapa Hifi Hotstart Ready Mix (KAPA Biosystems) with an amplicon PCR primer set (Forward: 5’-TCG TCG GCA GCG TCA GAT GTG TAT AAG AGA CAG CCT ACG GGN GGC WGC AG-3’, Reverse: 5’-GTC TCG TGG GCT CGG AGA TGT GTA TAA GAG ACA GGA CTA CHV GGG TAT CTA ATC C-3’). The PCR condition was: 95°C for 3 min, followed by 25 cycles of 95 °C for 30 sec, 55°C for 30 sec, and 72 °C for 30 sec, and then 72 °C for 5 min. PCR products were purified by AMpure XP beads (Beckman). Purified DNA was further amplified by PCR using Kapa Hifi Hotstart Ready Mix with Nextra XT Index Primers from Nextra XT Index Kit (Illumina). The PCR condition was: 95°C for 3 min, followed by 8 cycles of 95 °C for 30 sec, 55°C for 30 sec, and 72 °C for 30 sec, and then 72 °C for 5 min. After purification with AMpure XP beads, library DNA was quantified using a Qubit 1x dsDNA HS Assay Kit. Samples were sequenced on an Illumina Miseq with 2×300bp reads at the Sequencing Section at OIST

### 16S rRNA gene sequencing data analysis

FASTQ files were analyzed using the QIIME2 pipeline^65^ (QIIME2 version 2020.2). After conversion to the qza format, sequence data were demultiplexed and summarized using QIIME2 paired-end-demux. Then, sequences were trimmed and denoised with the dada2 plugin for QIIME2. Taxonomy was assigned using a naïve Bayes-fitted classifier trained on the SILVA_132 reference database (SSURef_NR99_132_SILVA) with the feature-classifier plugin for QIIME2. The phylogenetic tree for diversity analysis was reconstructed using QIIME2 align-to-tree-mafft-fasttree. Diversity analysis was performed with QIIME2 core-metrics-phylogenetic. PICRUSt2^66^ was used to determine predicted functions of bacterial communities. Comparisons of bacterial taxon abundance were performed with LEfSe^67^ using default parameters. In LEfSe analysis, reads assigned to the mitochondrial and chloroplast genomes were filtered out. In addition, taxa detected in less than 10% of participants (n = 86) or in less than 10% of a subset of participants (n = 40, top 20 and bottom 20 antibody or T cell responders) were excluded from the analysis.

### Treatment of PBMCs with SCFA and PGE2)

Cryopreserved PBMCs were thawed, centrifuged for 5 min at 440 g, and resuspended in TexMACS Medium (Miltenyi Biotec). PBMCs were seeded into a 96-well plate (10^6^ cells/well) and treated with SCFAs (mixture of 0.6 mM acetate (Sigma-Aldrich), 0.2 mM propionate (Sigma-Aldrich), and 0.2 mM butyrate (Sigma-Aldrich), 10 mM (*S*)-1, 2-Propanediol (Tokyo Chemical Industry), or 10 μM Prostaglandin E2 (Nacalai tesque). Cells were harvested at 18 h after treatment.

### RNA isolation and qPCR

cDNA was synthesized using ReverTra Ace qPCR RT Kit (Toyobo) using 200 ng of total RNA in a 10-μL volume. cDNA samples were diluted 4-fold by adding 30 μL sterile nuclease-free water and 10 μL of cDNA were used for PCR reactions. PCR was carried out using KAPA SYBR FAST qPCR Kit Master Mix (KAPA BIOSYSTEMS, KK4602) and primer sets (Supplementary Table 1) on a StepOnePlus Real-Time PCR System (Applied Biosystems).

### Statistical analysis

Statistical details for each experiment are included in the figure legends. Wilcoxon rank-sum tests and Wilcoxon signed-rank tests were performed using R (v.4.1.2) or GraphPad Prism (v.9.1.0). Correlation and partial correlation analyses were performed using Spearman’s correlation tests in the stats R package (v.4.1.2). For partial correlation tests, we removed the effects of age, gender, and fecal sampling timing from each dataset. P-values were corrected using Benjamini–Hochberg false discovery rate (FDR) for multiple comparisons.

## Data availability

The scRNA-seq, bulk RNA-seq, and 16S rRNA gene sequencing data that support the finding of this study have been deposited to DDBJ database under accession numbers DRA014613, DRA014614, and DRA014615, respectively. Any other relevant data are available from the corresponding author upon reasonable request.

## Acknowledgments

This study was in part supported by funding from COVID-19 AI and Simulation Project (Cabinet Secretariat) to HI, and the Platform Project for Supporting Drug Discovery and Life Science Research (BINDS) from AMED under grant number JP18am010107 to MW. We are also grateful to OIST Graduate University for its generous funding of the Immune Signal Unit. We thank physicians and nurses at KIN oncology Clinic and in Naha Medical association for excellent support to collect blood samples from donors and our laboratory members for valuable discussions. We also thank Steven D. Aird for editing the manuscript.

## Author contributions

MH performed data analysis of CyTOF, bulk and scRNA-seq, and 16S rRNA gene sequencing. MT performed most of the experiments except for ELISA and scRNA-seq. SY performed experiments and data analysis of ELISpot, CyTOF and 16S rRNA gene sequencing. NT prepared the scRNA-seq library and did some other experiments. M. Matthews and MW carried out anti-SARS-CoV-2 spike IgG ELISA. TT, YS, MY, ST, and Mio M. collected PBMCs from blood samples. TM, HT, OT, MK, ES, CY, Masataka M., and KT supervised volunteer recruitment and blood and stool sample collection. SY, MC, HK, and HI designed and supervised the study. MH, MT, and HI interpreted the data and wrote the manuscript.

## Conflicts of Interest

We declare that this research was conducted in the absence of any commercial or financial relationships that could be construed as a potential conflict of interest.

## Notes

### Competing Interest Statement

The authors have declared no competing interest.

