## Supplementary for "Human immune and gut microbial parameters associated with inter-individual variations in COVID-19 mRNA vaccine-induced immunity"

### Supplementary Figure 1

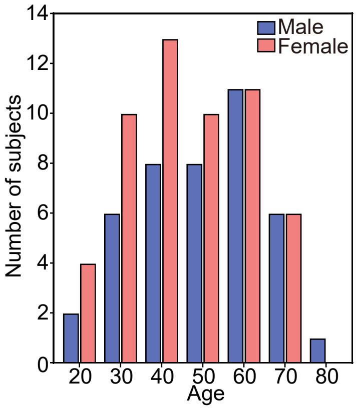


**Supplementary Fig. 1. Age and sex distribution of study participants.**
96 healthy subjects participated in this study.

### Supplementary Figure 2

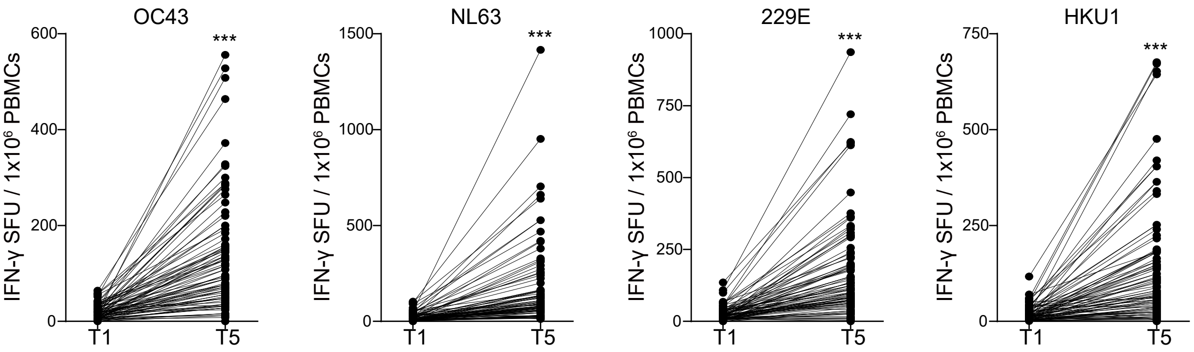


**Supplementary Fig. 2. BNT162b2-induced T-cell responses against human common cold coronaviruses.** IFN-γ-secreting T cells specific for spike of HCoV-OC43, NL63, 229E, and HKU1 in PBMCs at T1 and T5 were measured with ELISpot assays (n = 86). SFU, spot-forming unit. p values were calculated by Wilcoxon rank-sum tests (*** $p < 0.001$).

#### Supplementary Figure 3

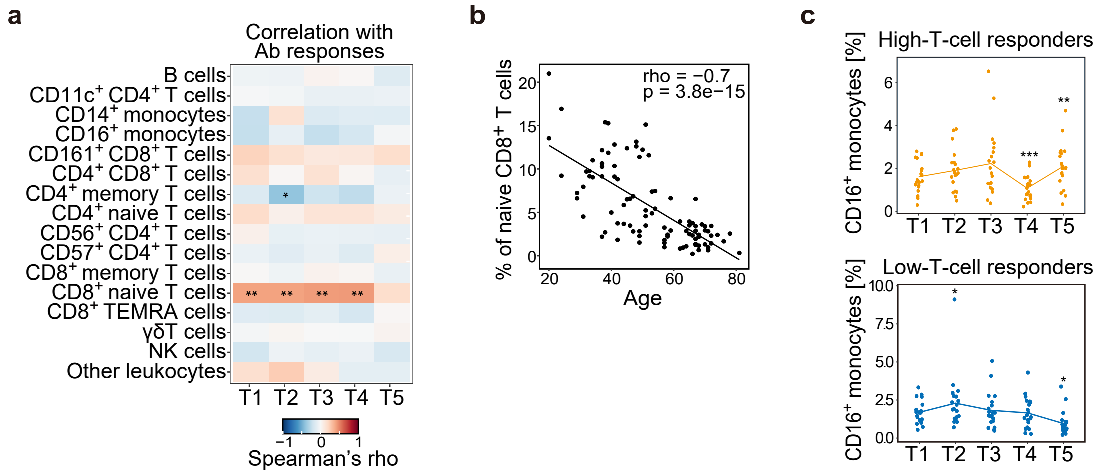


**Supplementary Fig. 3. The frequency of naive CD8$^+$ T cells is associated with BNT162b2-induced antibody responses. (a)** Heat map showing the correlation between the frequency of immune cell populations and BNT162b2-induced antibody responses. **(b)** Scatter plot showing a correlation between the baseline frequency of naive CD8$^+$ T cells and vaccine-induced antibody responses. **(a, b)** Correlation analyses without adjustments for age and sex were performed with Spearman's correlation tests with Benjamini–Hochberg FDR correction (* $p < 0.05$, ** $p < 0.01$). (c) Kinetics of the frequency of CD16$^+$ monocytes in PBMCs during vaccine response. High-T-cell responders (upper panel, n = 20) and low-T-cell responders (lower panel, n = 20) were analyzed. p values were calculated with Wilcoxon signed rank tests with Benjamini–Hochberg FDR correction (* $p < 0.05$, ** $p < 0.01$, *** $p < 0.001$).

#### Supplementary Figure 4

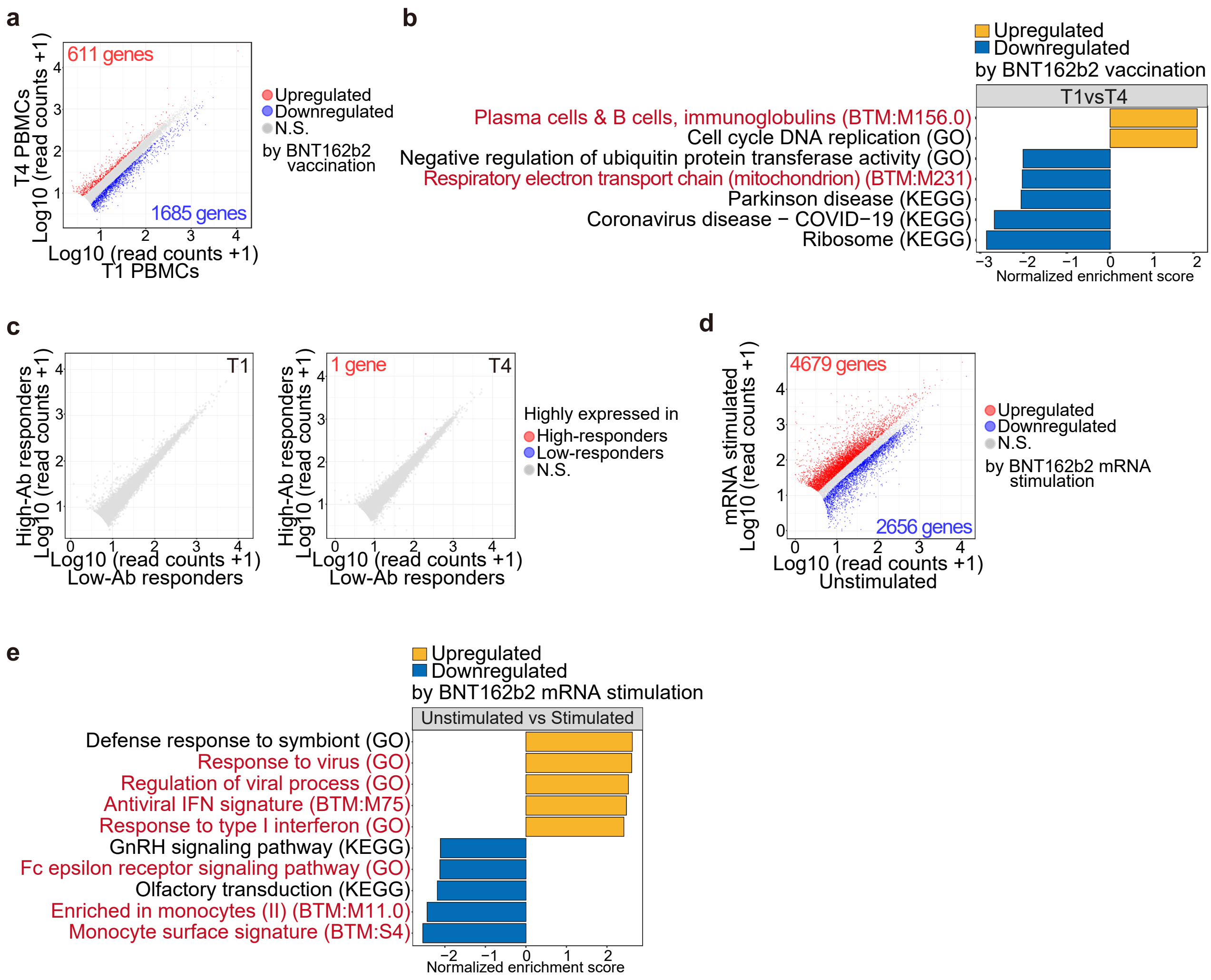


**Supplementary Fig. 4. BNT162b2-induced alteration of gene expression. (a-c)** RNA-seq data used in Fig. 4a-f were analyzed. **(a)** Scatter plots showing DEGs (log2 FC > 0.5, adjusted $p < 0.05$) between cells collected at T1 (n =80) and T4 (n = 78). Red and blue dots indicate DEGs that were upregulated and downregulated by BNT162b2 vaccination, respectively. N.S., not significant. **(b)** GSEA on a ranked gene list based on the fold change in expression between cells collected at T1 and T4. Immune-related BTM, GO, and KEGG pathways are shown in red. **(c)** Scatter plots showing DEGs (log2 FC > 0.5, adjusted $p < 0.05$) between high- (n = 18 at T1, n = 18 at T4) and low- (n = 19 at T1, n = 20 at T4) antibody responders. Red and blue dots indicate DEGs that were highly expressed in high- and low-antibody responders, respectively. **(d, e)** RNA-seq data used in Fig. 4g, h were analyzed. **(d)** Scatter plot showing DEGs (log2 FC > 0.5, adjusted $p < 0.05$) between cells unstimulated and stimulated with BNT162b2 mRNA for 6 h. Red and blue dots indicate DEGs that were upregulated and downregulated by BNT162b2 mRNA stimulation, respectively. **(e)** GSEA on a ranked gene list based on the fold change in expression between cells unstimulated and stimulated with BNT162b2 mRNA. Immune-related BTM, GO, and KEGG pathways are shown in red.

#### Supplemental Figure 5

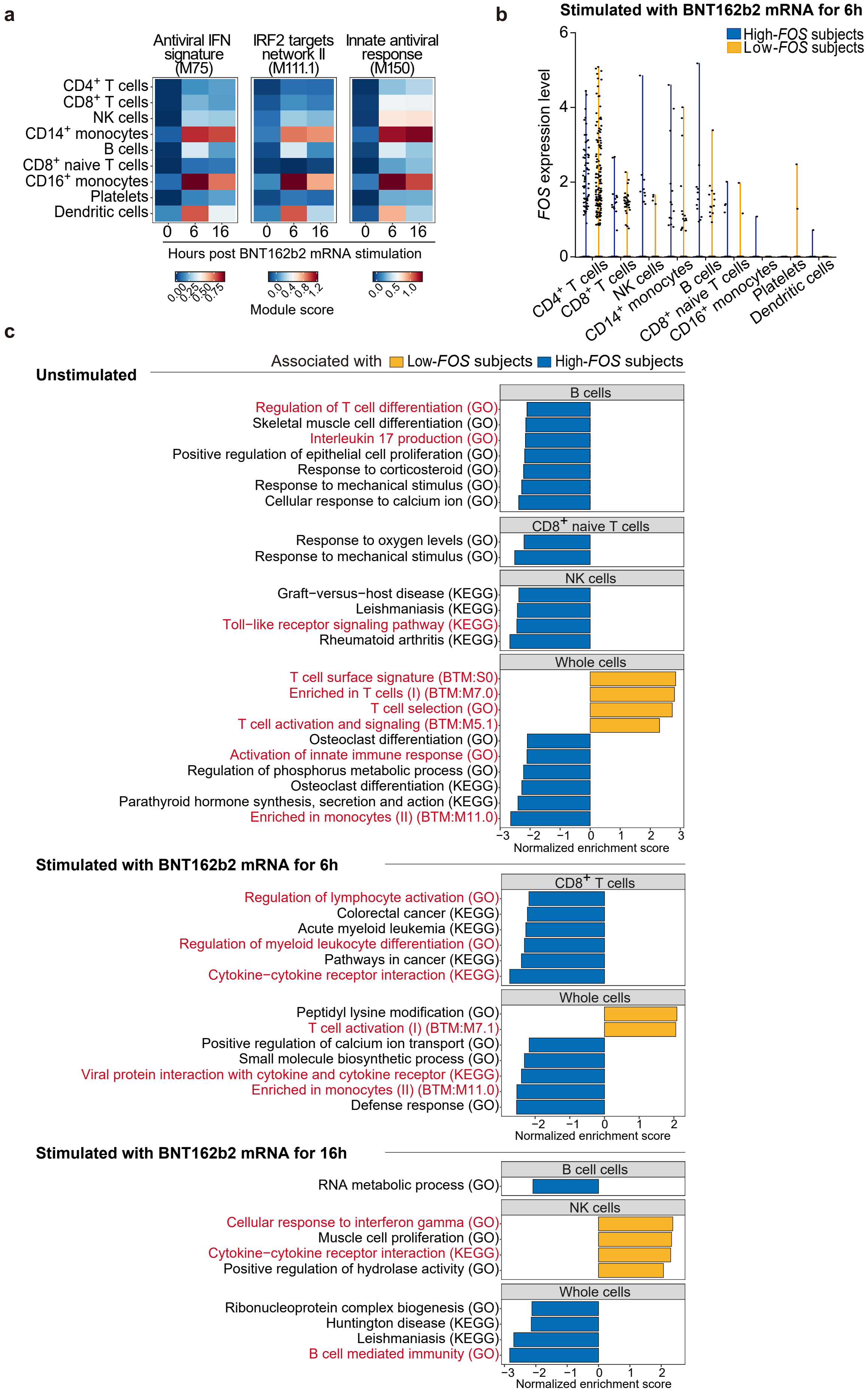


**Supplementary Fig. 5. Gene expression profiles in immune cell subpopulations unstimulated or stimulated with BNT162b2 mRNA.** scRNA-seq data obtained in Fig. 5 were analyzed. **(a)** Module score analysis of genes differentially expressed between unstimulated and stimulated immune cell populations. **(b)** Violin plots showing expression of *FOS* in PBMCs stimulated with BNT162b2 mRNA for 6 h (lower panel). **(c)** GSEA on a ranked gene list based on the fold change in expression in various immune cell populations unstimulated or stimulated with BNT162b2 mRNA for 6 or 16 h between high- and low-*FOS* subjects. Immune-related BTM, GO, and KEGG pathways are shown in red.

#### Supplementary Figure 6

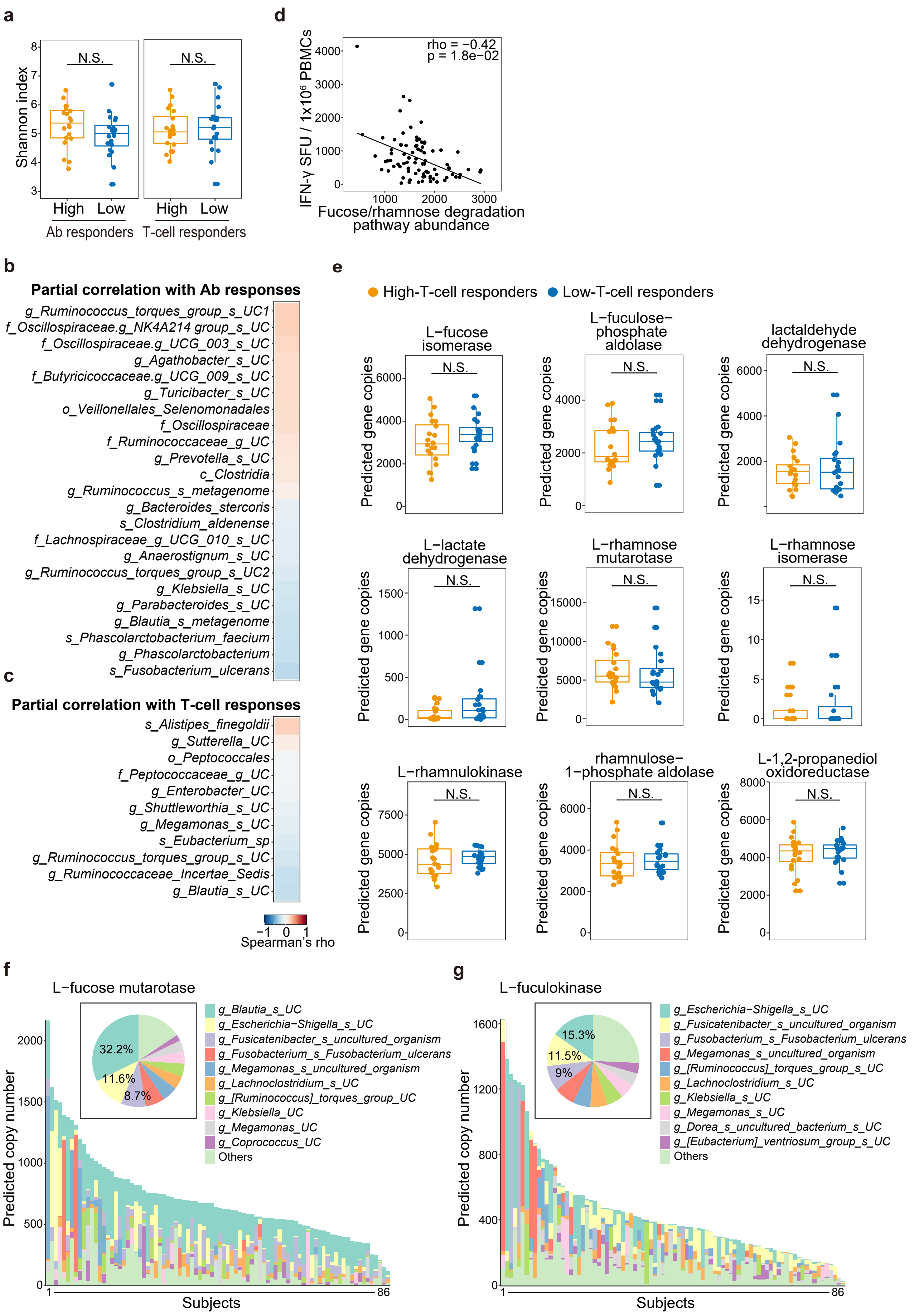

### Supplementary Figure 6

**Supplementary Fig. 6. Gut microbial taxa associated with BNT162b2-induced T cell responses.** 16S ribosomal RNA gene sequencing data obtained in Fig. 6 were analyzed. **(a)** Shannon index was compared in high vs low antibody (Ab) responders or in high- vs low-T-cell responders. p values were calculated with the Wilcoxon rank-sum tests with Benjamini–Hochberg FDR correction (N.S., not significant). **(b, c)** Heat map showing correlations between specific gut microbes and vaccine-induced antibody **(b)** or T-cell responses **(c)** (n = 86). Partial correlation analyses with adjustments for age, sex, and stool sampling timing were performed with Spearman's correlation tests with Benjamini–Hochberg FDR correction. There were no significant correlations between the bacterial taxa analyzed and vaccine-induced antibody or T-cell responses. o, order; f, family; g, genus; s, species; UC, unclassified. **(d)** Correlations of vaccine-induced T-cell responses with functions of gut microbiota were analyzed by Spearman's correlation tests with Benjamini–Hochberg FDR correction. Scatter plot showing an inverse correlation between the gut microbial fucose/rhamnose degradation pathway and vaccine-induced T-cell responses. **(e)** Analysis of the abundance of predicted gene copies for genes encoding enzymes involved in the fucose/rhamnose degradation pathway in high- and low-T-cell responders (n = 20 each). p values were calculated with the Wilcoxon rank-sum tests with Benjamini–Hochberg FDR correction (N.S., not significant). **(f, g)** The top 10 bacterial taxa that contributed to the abundance of predicted *L-fucose mutarotase* **(f)** and *L-fuculokinase* **(g)** genes were shown in pie charts (percentages in the whole subjects) and bar graphs (predicted gene copies in each subject).

**Supplementary Table1.** List of primer sets

| Name | Sequence |
|---|---|
| *hACTB*_F | ACAGAGCCTCGCCTTTG |
| *hACTB*_R | CCTTGCACATGCCGGAG |
| *hIFNb*_F | AAACTCATGAGCAGTCTGCA |
| *hIFNb*_R | AGGAGATCTTCAGTTTCGGAGG |
| *hCOX2*_F | CGGTGAAACTCTGGCTAGACAG |
| *hCOX2*_R | GCAAACCGTAGATGCTCAGGGA |
| *hFOS*_F | GCCTCTCTTACTACCACTCACC |
| *hFOS*_R | AGATGGCAGTGACCGTGGGAAT |
